# Monocyte-Mimetic Nanoprobe Enables Longitudinal MRI of Atherosclerotic Inflammatory Dynamics

**DOI:** 10.64898/2026.05.08.723851

**Authors:** Joshua Rousseau, Ting-Yun Wang, Sin-Pei Wu, Scott C. Beeman, Kuei-Chun Wang

## Abstract

Noninvasive monitoring of plaque inflammatory dynamics remains an unmet need. We previously developed a monocyte-mimetic nanoprobe, termed MoNP-SPION, for MRI detection of atherosclerotic lesions. Here we demonstrate MoNP-SPION enables longitudinal tracking of plaque inflammatory status in a clinically relevant mouse model. Following 16 weeks of plaque induction, mice were maintained on high-fat diet or switched to chow for 6 weeks to model persistent versus resolving plaque inflammation. MoNP-SPION-enhanced MRI was performed at 3- and 6-weeks post-adjustment, and arterial tissue was collected for histological assessment. Mice maintained on high-fat diet exhibited persistent hypointense T2* signal at the carotid bifurcation and aortic root, whereas chow-transitioned mice showed progressive signal attenuation, consistent with histological evidence of reduced plaque burden and inflammation. These findings establish MoNP-SPION as an effective molecular MRI probe for longitudinal assessment of plaque inflammatory dynamics, supporting its potential for monitoring atherosclerosis progression and therapeutic response.

## INTRODUCTION

Atherosclerosis, a leading cause of myocardial infarction and ischemic stroke, is a chronic inflammatory disease in which plaque composition and inflammatory status evolve as the disease progresses [1]. In response to cholesterol-lowering therapy, plaque burden may remain unchanged, whereas inflammatory activity can decline more substantially [2]. Thus, plaque inflammatory status may provide a more sensitive indicator of lesion activity and therapeutic response to assess plaque vulnerability. However, noninvasive assessment of plaque inflammatory dynamics remains an unmet clinical need, as conventional imaging modalities are largely anatomy-based and provide limited information on inflammation [3]. Molecular magnetic resonance imaging (MRI) offers a potential strategy to address this gap by enabling targeted visualization of inflammatory activity within vascular lesions.

Our previous work demonstrated that a monocyte-mimetic nanoprobe, which combines monocyte membrane-mediated targeting of inflamed vasculature with superparamagnetic iron oxide nanoparticle contrast (MoNP-SPION), enables sensitive MRI detection of atherosclerotic lesions [4]. Building on this, we extend the application of MoNP-SPION from single-timepoint lesion detection to longitudinal monitoring of plaque inflammatory status in a clinically relevant model of atherosclerosis. Specifically, MoNP-SPION-enhanced MRI was evaluated in atherosclerotic mice either maintained on high-fat diet (HFD) or switched to chow to model persistent versus resolving plaque inflammation. Our findings support MoNP-SPION as a molecular MRI probe capable of tracking plaque inflammatory dynamics over time, advancing its translational potential for assessing atherosclerosis progression and therapeutic response beyond anatomical changes.

## RESULTS

### MoNP-SPION-enhanced MRI detects longitudinal changes in vascular contrast over time

MoNP-SPION were formulated as previously reported (**Fig. 1A**) [4]. We employed a murine atherosclerosis model in which C57BL/6 mice were administered adeno-associated virus (AAV) encoding proprotein convertase subtilisin/kexin type 9 (PCSK9) to induce hypercholesterolemia and plaque formation during a 16-week HFD [5]. Following this induction period, mice were either maintained on HFD or switched to chow for an additional 6-weeks to model persistent hypercholesterolemia or dietary cholesterol lowering, respectively [2]. MoNP-SPION-enhanced MRI was performed at 3- and 6-weeks after diet adjustment in the carotid bifurcation and aortic root (**Figure 1B & C**). In the carotid bifurcation of mice maintained on HFD, post-injection images exhibited persistent hypointense blooming effects relative to pre-injection images. Quantification of normalized post-injection T2*-weighted signal intensities (SIs) yielded values of 79.84±5.97% and 74.39±9.33% at 3- and 6-weeks respectively (**Figure 1D**). In contrast, mice switched to chow showed reduced hypointense signal by week 3 and further resolution by week 6, resulting in post-injection normalized SIs of 88.97±3.62% and 91.71±4.91% respectively (**Figure 1D**). A similar trend was observed within the periluminal region of the aortic root. In mice maintained on HFD, hypointense signal remaining evident at both 3- and 6-weeks, with post-injection SIs of 77.35±4.02% and 79.29±3.14%, respectively (**Figure 1E**). Conversely, in mice adjusted to chow, periluminal hypointense delineation progressively diminished over the 6-week period, with post-injection SIs of 85.61±4.89% and 91.08±2.16% at 3- and 6-weeks, respectively (**Figure 1E**). Together, these data demonstrate that MoNP-SPION-enhanced MRI enables longitudinal monitoring of vascular T2* contrast dynamics in response to dietary intervention.

**Figure 1:**
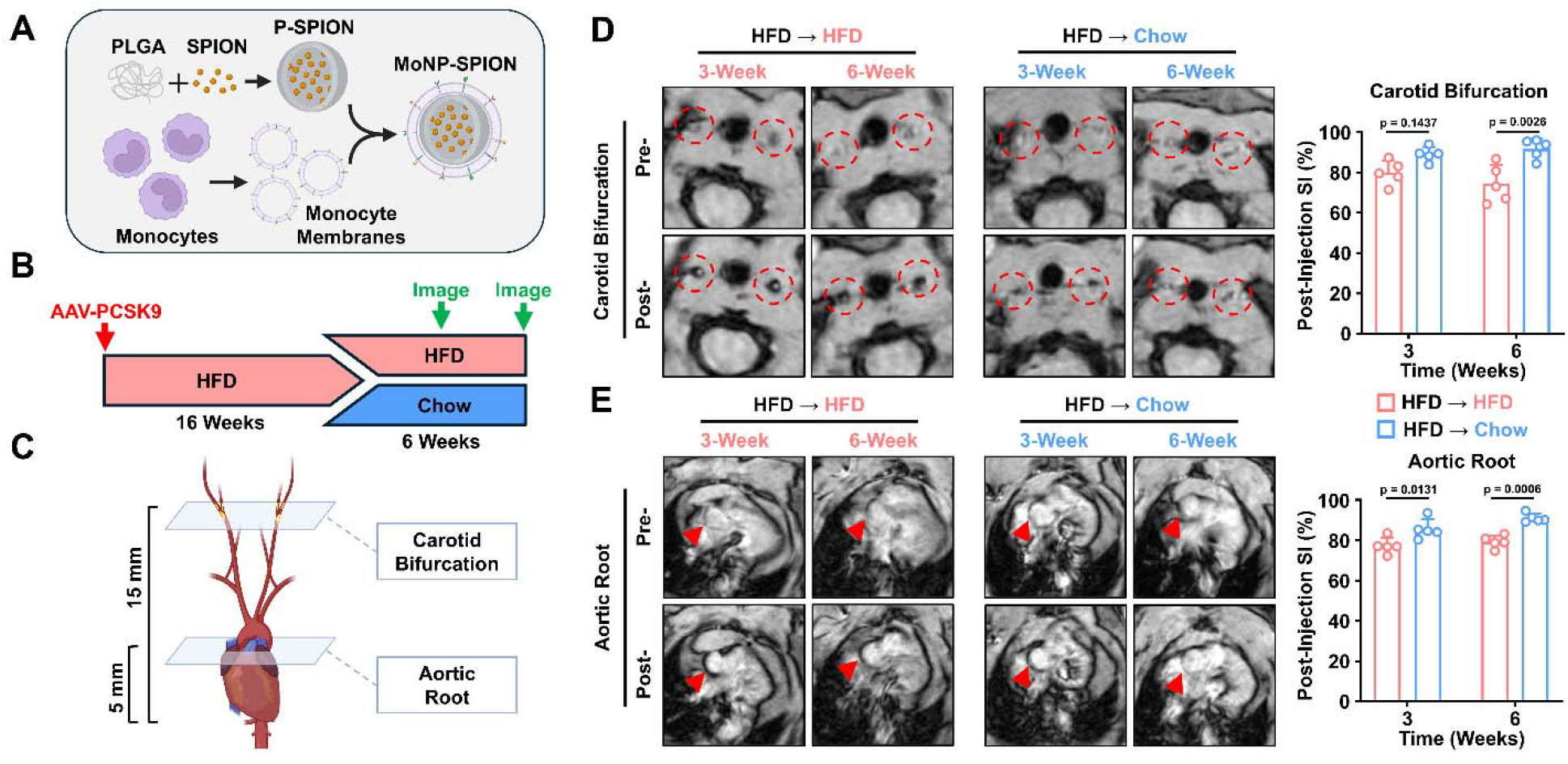
MoNP-SPION MRI detects diet-dependent changes in atherosclerotic plaques over time. (A-C) Schematic diagrams of (A) the experimental timeline, (B) imaging workflow, and (C) imaging planes for the carotid bifurcations and aortic root. (D, E) Representative pre- and post-MoNP-SPION T2^*^-weighted MRI images and post-injection normalized SI in the (D) carotid bifurcation and (E) aortic root of mice maintained on HFD or switched to chow. n = 5; two-way ANOVA with Tukey’s post hoc test.

### Imaging changes correspond to diet-dependent differences in plaque burden and macrophage infiltration

To validate the association between MoNP-SPION-induced contrast changes and plaque inflammatory status, we assessed systemic lipid levels, plaque burden, and macrophage infiltration after 6 weeks of dietary modulation. Mice maintained on HFD showed significantly higher total cholesterol levels than mice switched to chow (1512.20±348.40 mg/dL vs 331.60±74.88 mg/dL) (**Figure 2A**). Oil Red O staining further revealed significantly greater plaque burden in HFD mice, with lesions occupying an area of 0.12±0.03mm^2^ within the carotid bifurcation lumen and 0.68±0.13mm^2^in the aortic root lumen, compared with 0.07±0.01mm^2^ and 0.45±0.09mm^2^, respectively, in chow-fed mice (**Figures 2B & 2C**). Consistent with these differences, HFD mice exhibited greater CD68+ macrophage content in both the carotid artery and aortic root, whereas chow-fed mice showed markedly reduced macrophage infiltration (46.92±7.83% vs. 18.96±6.68% in carotid lesions; 64.76±8.82% vs. 35.80±2.99% in aortic root lesions) (**Figure 2D & 2E**). Together, these findings indicate that the persistent MoNP-SPION-induced T2* contrast observed in HFD mice corresponds to sustained plaque burden and macrophage-rich inflammation, whereas signal attenuation after chow transition reflects resolving inflammatory lesion status.

**Figure 2:**
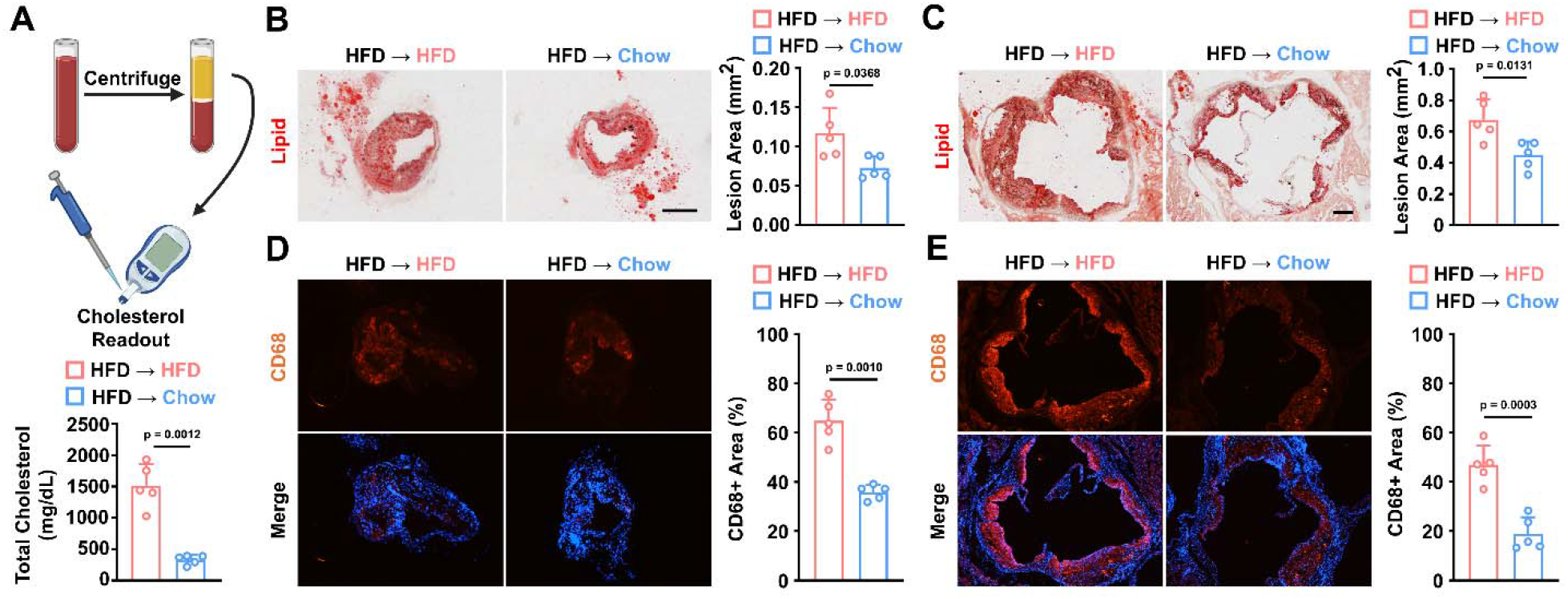
Dietary modulation alters plaque burden and inflammatory status. (A) Serum total cholesterol after 6-weeks of continued HFD or chow transition. (B, C) Oil red O-stained sections and lesion burden in (B) the carotid artery and (C) aortic root. (D, E) CD68 immunofluorescent staining and quantification in (D) the carotid artery and (E) aortic root. n = 5; unpaired t-test.

## DISCUSSION

Building on our previous demonstration that MoNP-SPION enhances MRI-based plaque detection compared with untargeted and free SPION formulations, the present study extends its application to longitudinal monitoring of plaque inflammatory dynamics. This capability is important because plaque inflammation is dynamic and closely associated with plaque vulnerability [6]. Molecular imaging with MoNP-SPION-enhanced MRI could therefore provide a more sensitive readout of plaque inflammatory activity, therapeutic response, and lesion risk than plaque burden alone.

A key strength of this study is the use of the PCSK9 gain-of-function model with dietary modulation, which provides a clinically relevant framework for evaluating inflammatory progression versus resolution in atherosclerosis. Consistent with previous studies, our histological analysis showed reduced plaque burden and CD68+ macrophage content in mice switched to chow, supporting plaque inflammatory resolution under reduced hypercholesterolemic condition [2]. Mechanistically, persistent T2^*^ hypointense signal in mice maintained on HFD is consistent with greater MoNP-SPION accumulation at inflamed vascular lesions, whereas signal attenuation after chow transition likely reflects reduced endothelial activation, diminished monocyte recruitment, and consequently lower MoNP-SPION accumulation [7]. The concordance between T2* signal changes and histological measures of plaque burden and macrophage content supports MoNP-SPION as an effective MRI probe of plaque inflammatory status.

Compared with positron emission tomography (PET)-based inflammation imaging, MoNP-SPION-enhanced MRI avoids ionizing radiation while providing high-resolution anatomical context and inflammatory status in the same imaging modality. Future studies evaluating MoNP-SPION in pharmacological lipid-lowering models and larger animal systems will further define its translational potential. Together, these findings demonstrate that MoNP-SPION not only identifies inflamed vasculature at a single time point, but also enables longitudinal assessment of plaque inflammatory dynamics, supporting its potential utility for monitoring atherosclerosis progression and therapeutic response beyond conventional anatomical imaging.

## MATERIALS AND METHODS

All animal experiments were approved by the Institutional Animal Care and Use Committee of Arizona State University (26-2173R). Eight-week-old male C57BL/6 mice received 5×10^11^ viral particles of AAV-PCSK9 (Vector Biolabs) via retro-orbital injection and placed on HFD (Inotiv, TD.88137) for 16 weeks, then either maintained on HFD or switched to chow for an additional 6 weeks [2,5]. MoNP-SPION were prepared and administered as previously described, and T2*-weighted MRI was performed at 3- and 6-weeks post-dietary modulation at the ASU-Barrow Center for Preclinical Imaging (TE: 3.5, TR: 600, α: 50) [4]. Following imaging, mice were euthanized, arterial tissues were sectioned and stained with Oil Red O for lipid content and CD68 antibody (Biolegend, #137001, 1:100 dilution) for macrophage infiltration and imaged on a Lionheart LX microscope. Serum was collected and lipid content was measured (Vetek Labs). Lesion and macrophage areas were quantified as percentages of luminal and lesion area respectively using ImageJ. All data are presented as mean ±SD (n=5). Statistical comparisons were performed using two-way ANOVA with Tukey’s post-hoc test or unpaired t-test in GraphPad Prism 10.1.2.

## ACKNOWLEDGEMENTS

This work was supported in part by R00135416 (to K.-C. W.). The authors acknowledge the use of the ASU-Barrow Center for Preclinical Imaging. Schematics created using BioRender.com.

## AUTHOR CONTRIBUTIONS

Conceptualization: J.R. and K.-C.W. Methodology: J.R., S.C.B., and K.-C.W. Investigation: J.R, T.-Y.W., and S.-P.W. Visualization: J.R. and K.-C.W. Supervision: K.-C.W. Writing – Original Draft: J.R. and K.-C.W.; Editing: J.R. and K.-C.W.

## COMPETING INTERESTS

The authors declare that they have no competing interests.

## Notes

### Competing Interest Statement

The authors have declared no competing interest.

### Funding

- National Institute of Health, USA: HL135416

